# 5’ cap sequestration is required for sensing of unspliced HIV-1 RNA by MDA5

**DOI:** 10.1101/2025.08.20.671346

**Authors:** Ivy K. Hughes, Siarhei Kharytonchyk, Sita Ramaswamy, Sallieu Jalloh, James B. Hood, Xianbao He, Andrew J. Henderson, Hisashi Akiyama, Alice Telesnitsky, Suryaram Gummuluru

## Abstract

Heterogenous transcription start site (TSS) usage dictates the structure and function of unspliced HIV-1 RNAs (usRNA). We and others have previously reported that expression and Rev/CRM1-mediated nuclear export of HIV-1 usRNA in macrophages activates MDA5, MAVS, and innate immune signaling cascades. In this study, we reveal that MDA5 sensing of viral usRNA is strictly determined by TSS, 5’ leader structure, and RNA function. We show that cap-sequestered HIV-1 usRNAs (^cap^1G) destined for viral genome packaging are specifically targeted by MDA5, while translation-destined (^cap^3G) usRNAs are remarkably immunologically silent. Using mutant viruses which express usRNA with altered 5’ cap-exposed leader structure, or inclusion of a retroviral constitutive transport element which drives mRNA-like NXF1-dependent nuclear export of viral usRNA, we show that cap exposure and nuclear export pathway choice are major determinants of both lentiviral RNA immunogenicity and function. In total, we identify innate immune system evasion as a possible rationale for the universal conservation of heterogenous TSS usage among ancestral and extant HIV-1 isolates and shed light on how MDA5 fundamentally discriminates between self and non-self RNAs.

**SIGNIFICANCE:** Innate immune activation is critical to both the process of initial infection establishment and ongoing chronic inflammation in HIV-1 infection. While MDA5 has been identified as the sensor which detects unspliced HIV-1 RNA produced in infected cells, it remains unclear how HIV-1 unspliced RNAs, which are generated by cellular transcriptional processes, are recognized as non-self. Here, we reveal that HIV-1 RNA function determines MDA5-driven immunogenicity. We show that only unspliced RNAs which traffic to membrane-associated viral assembly sites are immunogenic, while unspliced RNAs which are ribosomally translated to produce viral proteins are immunologically silent. These findings not only advance our knowledge of how the human innate immune system recognizes HIV-1 unspliced RNAs as foreign but also provide a rationale for the selective advantage to generate two pools of unspliced RNAs during HIV-1 replication.

## INTRODUCTION

The HIV-1 genome encodes a variety of viral transcripts across its ∼9 kb length whose expression is reliant on the promoter activity located at the 5’ long terminal repeat (LTR).(1) To accomplish this, HIV-1 employs a complex splicing strategy which allows for stoichiometric and temporal control over viral transcription.(2, 3) The first viral RNAs generated, encoding for Tat, Rev, and Nef proteins, are short, multiply spliced transcripts (msRNA) which are exported from the nucleus using canonical host mRNA export pathways dependent on NXF1.(4) Nuclear import of the *de novo*-translated Rev protein and subsequent engagement of the *cis*-acting Rev-response element (RRE) is associated with the generation of singly spliced (encoding *vif, vpr, vpu* and *env*) and full-length unspliced (usRNA, encoding *gag/pol* or acting as the viral genome) viral transcripts, which Rev links to the host factor CRM1 for nuclear export.(5, 6)

The relatively small HIV-1 single-stranded RNA genome encodes a remarkable amount of information not only in coding sequences across all three reading frames, but in structure as well. Highly structured regions in the full-length unspliced HIV-1 transcript such as those surrounding the *gag/pol* frameshift and RRE play documented roles in viral replication.(7–9) Among these regions is the 5’ untranslated region (5’ UTR), a structure-dense region of the HIV-1 genome which contains the *trans*-activation response element (TAR), dimerization initiation sequence (DIS), and major splice donor (SD) sequence, amongst others. Recent research has also revealed that the density of structural information encoded in the 5’ UTR is further enhanced by the transcription initiation promiscuity of host RNA polymerase II, which initiates differentially on a conserved GGG-tract at the 5’ end of the HIV-1 provirus and produces HIV-1 RNAs with either G-(^cap^1G) or triple GGG-(^cap^3G) 5’ end sequences.(10, 11) Remarkably, this 2-nucleotide difference at the 5’ end of usRNA leads to major changes in 5’ UTR structure and fate of HIV-1 usRNA.(12–14) In the ^cap^1G usRNA, the hairpin containing the polyA signal is intact, which leads to a 5’ UTR structure in which the 5’ cap structure is buried within the 5’UTR due to co-axial stacking of the TAR and polyA hairpins. In this conformation, the 5’ UTR DIS motif is exposed and available for dimerization with a second gRNA for packaging into nascent particles.(15) This contrasts with the ^cap^3G usRNA, in which the polyA hairpin is unstable and instead forms a more continuous central hairpin-like structure which includes DIS, sequestering the signals needed for packaging and instead exposing the 5’ cap needed for interactions with translation initiation factors and ribosomal translation.(16, 17)

We and others have previously reported that cytoplasmic expression of intron-containing HIV-1 RNAs exported from the nucleus via Rev/CRM1 activate innate immune responses in macrophages and dendritic cells in an MDA5-dependent manner.(18–21) MDA5, a RIG-I like receptor (RLR) family member, surveys the cytoplasm for viral or host-derived immunostimulatory RNAs, binding of which induces oligomerization of MDA5 through N-terminal caspase activation and recruitment domains (CARD) and subsequent recruitment and activation of MAVS through CARD-CARD interactions. We have recently demonstrated that activation of downstream TRAF6 E3 ubiquitin ligase, IKK-β kinase, and IRF5 are required for expression of type I interferons (type I IFN) and proinflammatory cytokines in HIV-1-infected macrophages.(21) While the host nucleic acid sensing and signal transduction machinery that detects HIV-1 intron-containing RNAs and activates type I IFN responses has been elucidated, sequence and structural requirements of the viral transcripts that define their immunostimulatory potential have remained unclear. In this report, using primary human monocyte-derived macrophages as a model of myeloid cell infection, we show that innate immune responses to intron-containing HIV-1 RNA initiated by MDA5 are restricted to ^cap^1G usRNA specifically. Further, we show that MDA5-mediated immunogenicity is linked to nuclear export pathway, cytoplasmic RNA fate, and 5’ cap exposure, implying that HIV-1 ^cap^3G usRNAs destined for translation make use of host RNA metabolism pathways to dampen immunogenicity while facilitating viral protein expression. In summary, this work provides a new model to understand the ability of MDA5 to distinguish 5’ UTR structures of HIV-1 usRNAs based on 5’ cap exposure and the functional importance of TSS heterogeneity in the context of innate immune activation by HIV-1.

## RESULTS

### *De novo*-transcribed ^cap^1G HIV-1 unspliced RNAs stimulate innate immunity in macrophages

In the HIV-1 LTR, a highly conserved GGG motif encompasses the transcription initiation site and is located approximately 30 bp downstream of the viral core promoter CATA box element.(22) The LTR sequence in the HIV-1/NL4-3 lab-adapted strain is sufficient to drive transcription of both ^cap^1G and ^cap^3G usRNAs (**Fig. 1A/B**).(10, 12) Mutation of the first two G residues in the NL4-3 LTR to TC instead drives transcription only from the final G in the initiation tract, producing a virus in which only ^cap^1G RNA is transcribed (1G virus, where 1G refers to the mutant virus and ^cap^1G refers to the RNA transcripts it produces). Similarly, if the first two G residues of the GGG tract are not replaced but instead shifted 2 bp downstream by insertion of the TC motif, transcription results in only ^cap^3G RNA expression (3G virus) (**Fig. 1B**). To determine the immunological outcome of ^cap^1G and ^cap^3G HIV-1 usRNA expression in macrophages, we infected primary human monocyte-derived macrophages (MDMs) with VSV-G-pseudotyped replication-competent viruses (containing both G and Env surface expression). To overcome SAMHD1-mediated inhibition of reverse transcription in primary MDMs, we co-infected with SIVmac Vpx-containing VLPs (SIV3+ VLPs) as previously described.(23) We first ensured that 1G and 3G infection of MDMs resulted in the specific transcription of only ^cap^1G and ^cap^3G RNAs by harvesting RNA from infected macrophages and performing a cap-dependent adaptor ligation/PCR assay (CaDAL) for viral usRNA as previously described.(22) Indeed, we observed highly specific expression of unspliced ^cap^1G/3G RNAs upon infection with 1G/3G mutant viruses, respectively (**Fig. 1C**). Further, staining for intracellular p24^Gag^ by flow cytometry and measuring p24^Gag^ release in culture supernatants by ELISA at three days post infection confirmed that the efficiency was equivalent between WT, 1G, and 3G virus infections of MDMs (**Fig. 1D/E**).

**Fig. 1:**
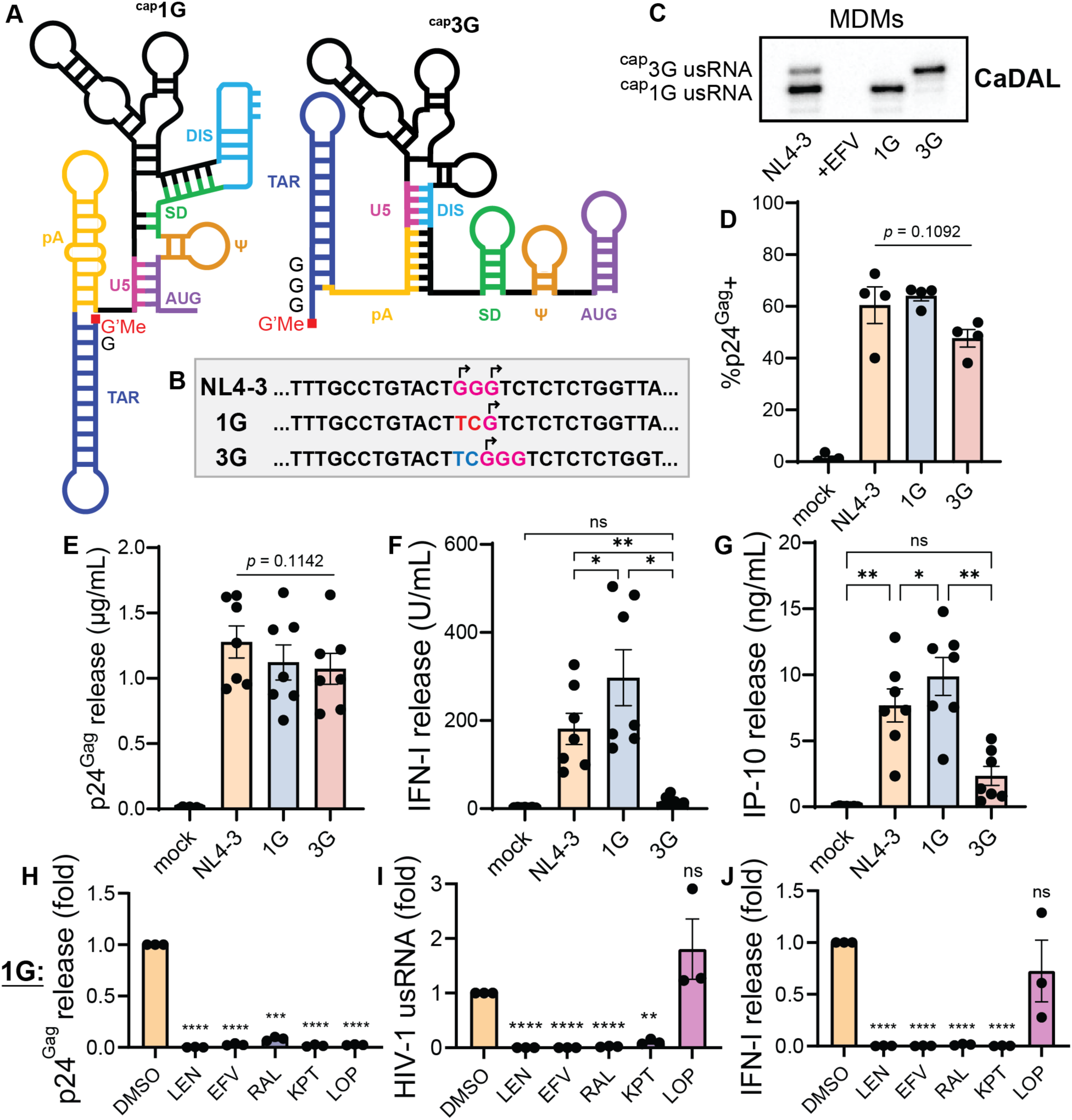
*De novo*-transcribed ^cap^1G HIV-1 unspliced RNAs stimulate innate immune responses in macrophages. (**A**) Schematic of 2D RNA structures demonstrating the predominant folded 5’ leader conformations of ^cap^1G and ^cap^3G unspliced HIV-1 RNAs. (**B**) 5’ proviral sequences of wild-type NL4-3, ^cap^1G-only-expressing NL4-3 (1G), and ^cap^3G-only-expressing NL4-3 (3G) constructs. The GGG transcription initiation tract is highlighted in pink, with ^cap^1G and ^cap^3G initiation sites labeled with arrows. In 1G virus, GG is mutated to TC (in red). In 3G virus, TC is inserted (in blue) which pushes the GGG initiation tract 2 nt downstream. (**C**) **Ca**p-**d**ependent **a**daptor **l**igation PCR assay (**CaDAL**) analyzing 5’ ends of unspliced HIV-1 RNA expressed in virus-infected MDMs. +EFV refers to NL4-3 infection with efavirenz treatment. (**D**-**I**) Primary human MDMs were co-infected with Vpx-containing SIV3+ VLPs and VSV-G-pseudotyped HIV-1/NL4-3. Plots show (**D**) infection efficiency evidenced by p24^Gag^ immunostaining analyzed by flow cytometry, (**E**) p24^Gag^ secretion into the culture supernatant assayed by ELISA, (**F**) bioactive type I IFN secretion assayed by bioassay, (**G**) IP-10 secretion assayed by ELISA, all at 3 dpi. (**H**-**J**) MDMs were co-infected with SIV3+ VLPs and 1G virus in the presence of antiretrovirals lenacapavir (capsid, 1 µM), efavirenz (reverse transcriptase, 1 µM), raltegravir (integrase, 30 µM), KPT-330 (Rev/CRM1, 1 µM), or lopinavir (protease, 1 µM). Plots show relative p24^Gag^ release (**H**, specific for cleaved p24^Gag^), HIV-1 usRNA expression by RT-qPCR (**I**), and IFN-I release (**J**) at 3 dpi. (**D**-**J**) Data was analyzed by one-way ANOVA with Tukey’s post-test (**D**-**G**) or Dunnett’s post-test with statistics referring to comparison with DMSO condition (**H**-**J**). (**D**/**E**) All comparisons within virus infection conditions are nonsignificant. (**D**-**J**) Each data point is a primary cell donor with plots showing mean +/- SEM, ns: p≥0.05; *: p<0.05; **: p<0.01; ***: p<0.001; ****: p<0.0001.

To measure type I IFN secretion, we employed a bioassay to measure total bioactive type I IFN levels. Though infection establishment was equivalent across all three viruses, we observed type I IFN secretion only from NL4-3- and 1G-infected MDMs, but not from 3G-infected MDMs (**Fig. 1F**). IFN-I secretion from 1G-infected MDMs was also significantly greater than that observed with wild-type NL4-3-infected MDMs, consistent with specific sensing of ^cap^1G RNAs. We also observed significantly less pro-inflammatory IP-10 secretion from 3G-infected cultures compared to 1G or wild type virus infected MDMs (**Fig. 1G**). We confirmed that innate immune activation was limited to *de novo*-transcribed ^cap^1G usRNA rather than incoming virus-associated gRNAs since inhibition of infection establishment (lenacapavir, efavirenz, and raltegravir) or Rev/CRM1 usRNA export (KPT-330), but not protease activity (lopinavir) blocked induction of type I IFN secretion (**Fig. 1H**-**J**). These findings are consistent with the established model that innate immune activation is induced late in the virus lifecycle after successful Rev/CRM1-mediated export of HIV-1 usRNA.(18–21) Additionally, we infected MDMs with NL4-3, 1G, and 3G viruses without addition of Vpx-containing SIV3+ VLPs, and observed type I IFN secretion peaking at 6 days post-infection in wild-type and 1G infections but not 3G infection despite identical levels of *de novo* p55^Gag^ expression and p24^Gag^ secretion (**Fig. S1**). Together, these findings establish that expression of unspliced ^cap^1G RNAs alone stimulates proinflammatory type I IFN responses in macrophages.

### MDA5 detects ^cap^1G unspliced HIV-1 RNAs and activates type I interferon expression through MAVS

We and others have previously reported that MDA5 senses HIV-1 usRNAs and activates MAVS and downstream innate immune responses.(20, 21) To determine whether MDA5 detects ^cap^1G usRNA, we transfected MDMs with siRNAs targeting MDA5, RIG-I, or MAVS and infected cells with wild-type (1G/3G) or 1G viruses and measured induction of innate immune responses. We ensured that targeted siRNA transfection resulted in efficient knockdown of MDA5, RIG-I, and MAVS expression (**Fig. 2A**-**F**). We also ensured that siRNA transfection did not impact infection efficiency or expression of usRNA, evidenced by equivalent levels of p24^Gag^ secretion and usRNA quantification via RT-qPCR in siRNA-transfected MDMs (**Fig. 2G/H**). We did not observe any significant diminution of cytokine secretion in siRIG-I-transfected wild-type or 1G virus infected MDMs (**Fig. 2I/J**). Consistent with our hypothesis, type I IFN and IP-10 release were abrogated in siMDA5- and siMAVS-transfected MDMs infected with wild-type or 1G virus.

**Fig. 2:**
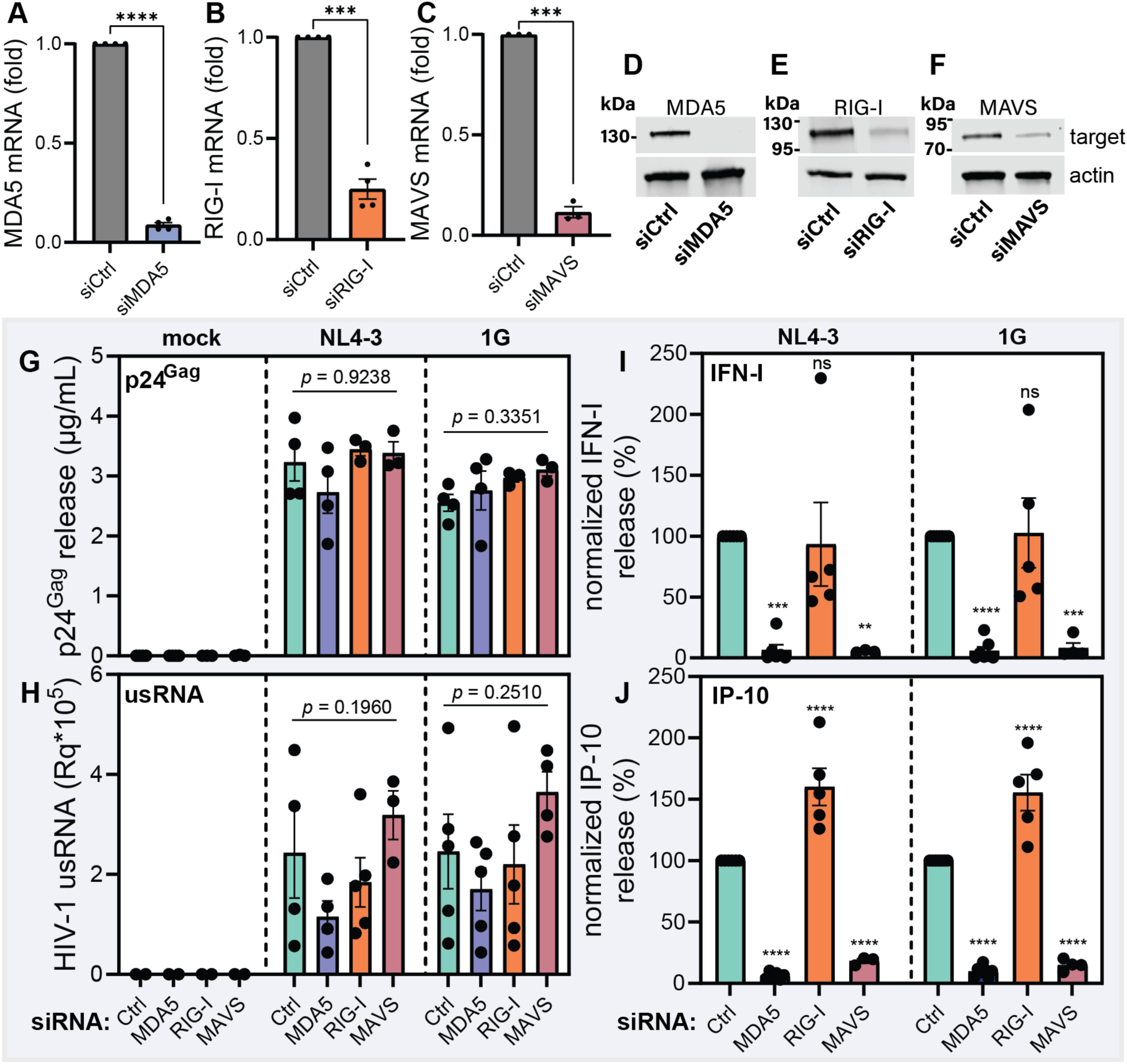
MDA5/MAVS expression is required for ^cap^1G HIV-1 usRNA-induced innate immune responses in macrophages. MDMs were transfected with siRNA targeting the indicated sensors/adaptor and two days later co-infected with SIV3+ VLPs and VSV-G-pseudotyped HIV-1/NL4-3. (**A**-**C**) RT-qPCR of RNA extracted from uninfected siRNA-transfected MDMs normalized to GAPDH expression. (**D**-**F**) Western blots of uninfected siRNA-transfected MDMs probed with the indicated antibodies. (**G**/**H**) p24^Gag^ release (**G**) and RT-qPCR for HIV-1 usRNA normalized to GAPDH (**H**) at 3 dpi. (**I**/**J**) type I IFN release (**I**) and IP-10 release (**J**) normalized to siCtrl-transfected infected MDMs. (**D**-**F**) Western blots are of one representative primary cell donor. (**A**-**C**,**G**-**J**) Data points are individual primary cell donors. Plots show mean +/- SEM analyzed by paired t-test (**A**-**C**) or two-way ANOVA with Tukey’s post-test (**G**/**H**) or Dunnett’s post-test with statistics referring to siCtrl-transfected condition (**I**/**J**), ns: p≥0.05; **: p<0.01; ***: p<0.001; ****: p<0.0001. (**G**/**H**) all comparisons within the indicated virus infection condition are nonsignificant.

To confirm that MDA5 was responsible for specifically engaging viral ^cap^1G usRNA transcripts, we analyzed the *in situ* interaction of MDA5 with unspliced viral transcripts by performing RNA immunoprecipitation in virus-infected HEK293T cells overexpressing MDA5-FLAG (**Fig. 3A**).(21) Following UV crosslinking and anti-FLAG immunoprecipitation, we observed significant infection-dependent anti-FLAG pulldown of unspliced HIV-1 transcripts in NL4-3 and 1G virus infection conditions relative to control IgG pulldown, but no usRNA pulldown in 3G-infected MDA5-FLAG-expressing HEK293T cells (**Fig. 3B**). Consistent with our hypothesis, we did not observe pulldown of multiply-spliced viral transcripts or host actin mRNA in any virus infection condition (**Fig. 3C/D**). Taken together, these studies indicate that MDA5 senses productive HIV-1 infection by specific interaction with newly-transcribed ^cap^1G unspliced transcripts, leading to MAVS activation and induction of innate immune responses in macrophages.

**Fig. 3:**
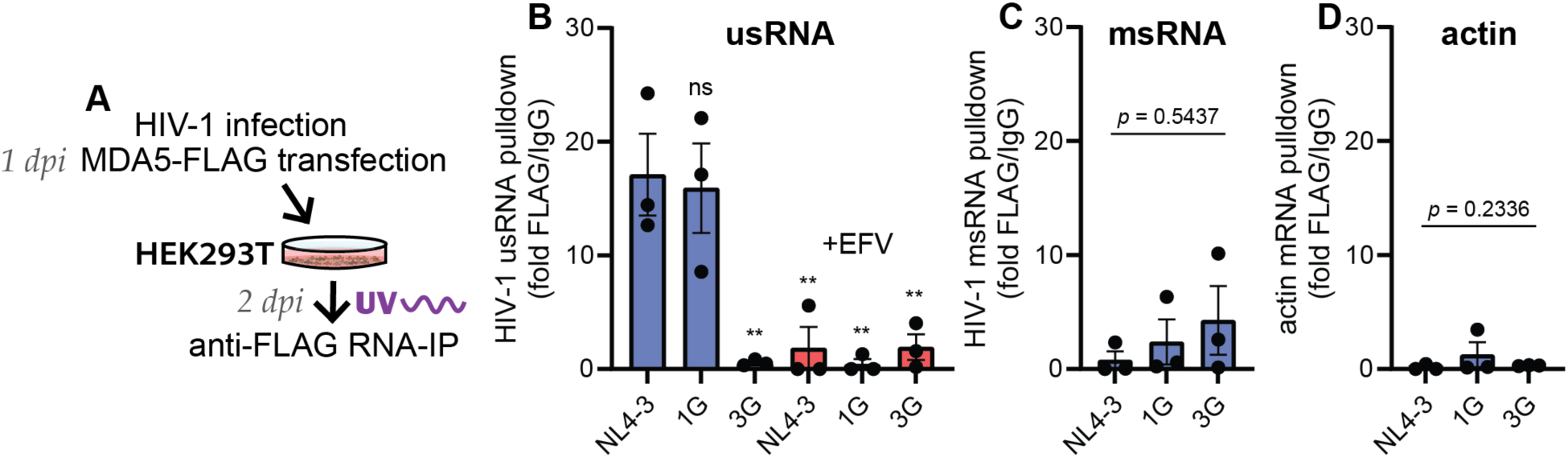
MDA5 specifically engages ^cap^1G HIV-1 usRNA. (**A**) HEK293T cells were infected with VSV-G-pseudotyped replication-competent viruses in the presence or absence of 1 µM efavirenz. Infected cells were transfected with MDA5-FLAG plasmid. Cells were UV-crosslinked and cytosolic fractions underwent RNA immunoprecipitation at 2 dpi using anti-FLAG or IgG control antibodies. (**B**-**D**) Plots show RT-qPCR fold enrichment of MDA5-FLAG signal over IgG control antibody conditions for (**B**) HIV-1 unspliced RNA, (**C**) HIV-1 multiply spliced RNA, and (**D**) actin mRNA. Plots show mean +/- SEM and each data point is an independent experiment. (**B**) One-way ANOVA with Dunnett’s post-test was used with statistics referring to comparison with NL4-3 condition, ns: p≥0.05; **: p<0.01. (**C**/**D**) One-way ANOVA with Tukey’s post-test was used, all comparisons nonsignificant.

### ^cap^1G/3G RNA splicing and usRNA localization in primary macrophages

To better understand the dynamics of MDA5-driven innate responses in HIV-infected cells, we characterized ^cap^1G RNA metabolism and fate in primary macrophages. We analyzed ^cap^1G and ^cap^3G unspliced RNA abundance in wild type (^cap^1G/3G) virus-infected syngeneic CD4+ T cells and macrophages using CaDAL assay. In CD4+ T cells, we found a roughly 1:1 ratio of *de novo*-transcribed ^cap^1G/^cap^3G unspliced RNA species in infected cell lysates, which was absent in efavirenz-treated cell lysates (**Fig. 4A**). In MDMs, we observed a distinct shift towards ^cap^1G unspliced RNA abundance which was again dependent on establishment of productive infection (**Fig. 4A**). We next analyzed usRNA levels in the wild-type, 1G, or 3G virus infections of MDMs by RT-qPCR, and again observed a decrease in unspliced HIV-1 RNA signal in 3G infection compared to 1G and wild-type NL4-3 infection conditions (**Fig. 4B**). We hypothesized that the low usRNA abundance in 3G-infected MDMs was either due to increased splicing of ^cap^3G RNA products, or a shorter cellular half-life of ^cap^3G usRNA, or both. To test these hypotheses, we analyzed the viral RNA transcriptome in MDMs using a multiplex digital-droplet PCR (ddPCR)-based assay, simultaneously analyzing the presence of unspliced (LTR/gag/env/nef), singly-spliced (LTR/env/nef), or multiply-spliced (LTR/nef) transcript sequences using previously defined primer/probe sets.(24) We observed a significant shift towards unspliced RNA fate (positive for all four probes simultaneously) in 1G infection compared to NL4-3 and 3G infection of MDMs (**Fig. 4C**, **Fig. S2**). This effect was accompanied by a decrease in singly-spliced RNA in 1G infection, and a roughly equivalent abundance of multiply-spliced RNA in all three infections, similar to findings in the previous report.(25)

**Fig. 4:**
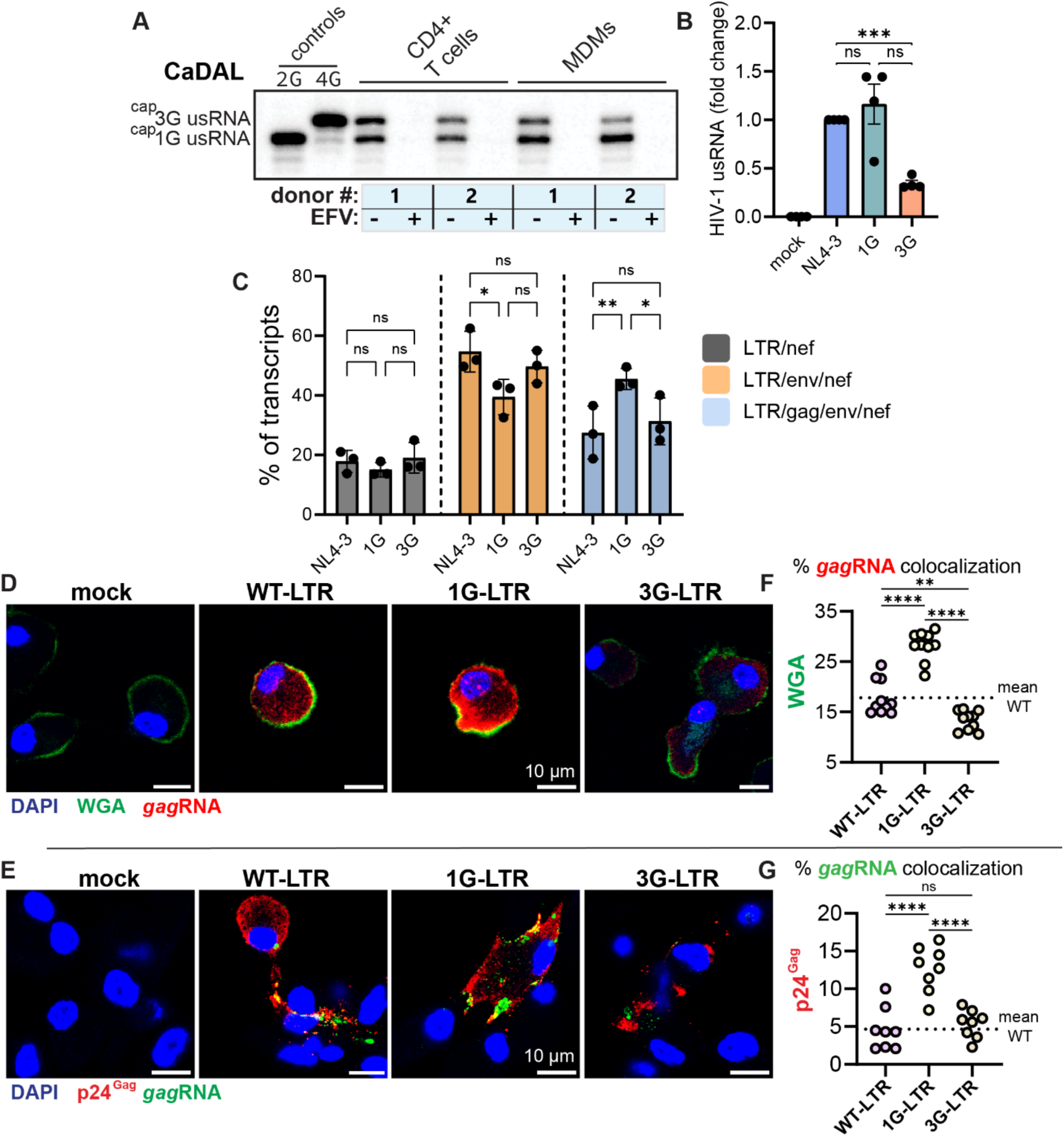
Transcription and intracellular localization of ^cap^1G and ^cap^3G usRNA in macrophages. (**A**) CaDAL assay analyzing unspliced HIV-1 transcripts using RNA isolated from syngeneic activated primary CD4+ T cells and MDMs infected with replication-competent VSV-G-pseudotyped NL4-3 (with SIV3+ VLP co-infection for MDMs) with or without efavirenz pre-treatment. Comparisons within samples are quantitative, comparisons between samples are not quantitative. (**B**) Relative RT-qPCR for unspliced HIV-1 transcripts using RNA isolated from MDMs co-infected with HIV-1/SIV3+ VLPs. (**C**) ddPCR transcriptomic analysis of RNA used in (**B**). cDNA was generated using random hexamers and analyzed with multiplex ddPCR analyzing LTR, *gag*, *env*, and *nef* expression. Droplets containing single cDNA molecules were classed as multiply-spliced (LTR/nef), singly-spliced (LTR/env/nef), or unspliced (LTR/gag/env/nef) or discarded (other combinations). Transcript abundance was normalized for each donor (all three populations totaling 100% for each donor). (**D**-**F**) MDMs were co-infected with SIV3+ VLPs and single-round (Δenv/GFP) viruses harboring wild-type, 1G, or 3G LTRs. smRNA-FISH was performed targeting unspliced *gag* RNA with WGA (**D**, wheat germ agglutinin, plasma membrane stain) or anti-p24^Gag^ co-stain (**E**). (**F**/**G**) Plots show % colocalization of *gag*RNA signal with WGA or p24^Gag^. (**A**) CaDAL was performed with two primary cell donors. (**B**/**C**) Data was analyzed with one-way or two-way ANOVA with Tukey’s post-test. Data is shown as mean +/- SEM, all data points are individual primary cell donors, ns: p≥0.05; *: p<0.05; **: p<0.01; ***: p<0.001. (**D**/**E**) Images show one representative field. (**F**/**G**) Data points are individual fields of view within one experiment, representative of over three independent experiments. Data was analyzed using one-way ANOVA with Tukey’s post-test, ns: p≥0.05; **: p<0.01; ****: p<0.0001.

To characterize the localization of ^cap^1G/3G usRNA expression in primary macrophages, we employed single-molecule RNA-FISH (smFISH) with co-staining for plasma membrane (wheat germ agglutinin, WGA) or p24^Gag^ expression (**Fig. 4**D/E, **Fig. S3**). For these experiments, we employed single-round (Δenv, GFP in place of *nef*) viruses harboring wild-type (^cap^1G/3G, WT-LTR) or ^cap^1G-or ^cap^3G-only LTRs (1G-LTR, 3G-LTR). We also ensured that cytoplasmic usRNA signal was dependent on productive infection using reverse transcriptase inhibition (**Fig. S3**), and Rev/CRM1 nuclear export using a virus with an inactivating mutation in Rev (WT-LTR/M10, **Fig. S3**).(18, 26) usRNA expressed in 1G infection of MDMs preferentially co-localized with both WGA-positive plasma membrane and sites of Gag accumulation (**Fig. 4F/G**), consistent with the established function of ^cap^1G usRNA transcripts as packaging-fated virus genomes.(12, 27) Taken together, these experiments suggest that HIV-1 ^cap^1G usRNA accumulate to higher levels at membrane-associated sites of viral assembly in primary macrophages, and that preferential accumulation at these peripheral membrane sites may contribute to specific surveillance of ^cap^1G usRNA by MDA5 and induction of innate immune responses.

### RNA fate links expression of ^cap^1G unspliced HIV-1 RNA to innate immune activation

While both unspliced ^cap^1G and ^cap^3G contain highly structured 5’ UTRs, ^cap^1G unspliced RNAs predominantly dimerize to serve as viral genomes, which could theoretically explain their susceptibility to MDA5 sensing. To test if dimerization is the trigger of innate immune responses in macrophages, we infected MDMs with an additional mutant virus which results in the expression of usRNA with a novel 5’ leader structure. This virus construct, known as no bulge (NB), encodes a polyA hairpin with greater internal base-pairing and stability (**Fig. 5A**), which when expressed from a ^cap^3G-only LTR (3G-NB virus) produces a single ‘hybrid’ unspliced RNA (in addition to typical spliced transcripts) in which the 5’ cap is exposed as in typical ^cap^3G usRNA, but which contains an overall ^cap^1G-like fold characterized by a double three-way junction structure and exposure of DIS.(28) Previous work has established that while ^cap^3G-NB usRNAs dimerize *in vitro*, they are efficiently translated in cells and not packaged in competition assays with ^cap^1G usRNA, suggesting that 5’ cap exposure is the major determinant of whether an usRNA will adopt a packaging or translation fate.(27, 28) When we infected primary MDMs with WT, 3G, and 3G-NB viruses, we observed similar levels of infection efficiency and p24^Gag^ release for all viruses (**Fig. 5B**/**C**, **Fig. S4**).

**Fig. 5:**
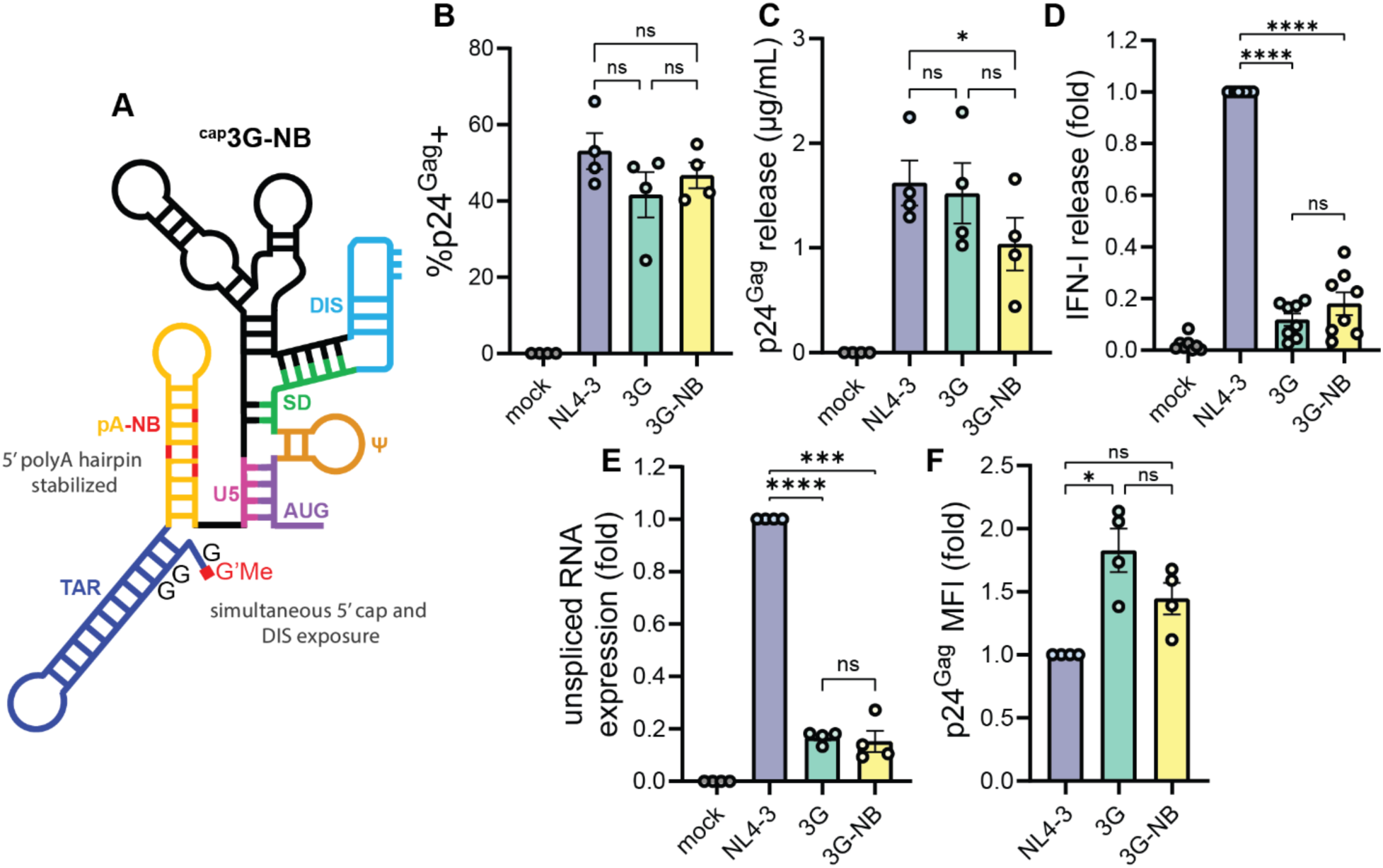
5’ cap sequestration on HIV-1 usRNA determines MDA5-driven immunogenicity. (**A**) Schematic of the predominant folded 5’ leader structure of ^cap^3G unspliced HIV-1 RNA with ‘no bulge’ (NB) contact-stabilizing mutations in the polyA hairpin. (**B**-**F**) MDMs were co-infected with replication-competent VSV-G-pseudotyped HIV-1/SIV3+ VLPs. Plots show (at 3 dpi) (**B**) infection efficiency analyzed by flow cytometry after anti-p24^Gag^ antibody staining, (**C**) p24^Gag^ release into the culture medium, (**D**) relative bioactive type I IFN secretion into the culture medium, (**E**) relative RT-qPCR for HIV-1 usRNA normalized to GAPDH expression, and (**F**) p24^Gag^ flow cytometry mean fluorescence intensity (MFI) of p24^Gag^-positive cells. Plots show mean +/- SEM and each data point is an individual primary cell donor. Data was analyzed by one-way ANOVA with Tukey’s post-test, ns: p≥0.05; *: p<0.05; **: p<0.01; ***: p<0.001; ****: p<0.0001.

When we measured bioactive type I IFN from these cultures, we found that 3G-NB virus infection resulted in minimal IFN secretion, similar to that observed upon 3G infection (**Fig. 5D**). Interestingly, relatively low abundance of usRNA transcripts in 3G-NB infection was observed, similar to 3G infection of MDMs (**Fig. 5E**). However, despite these low usRNA levels, we observed high levels of Gag expression in 3G- and 3G-NB-infected MDMs (**Fig. 5F**, **Fig. S4**), suggesting that 5’ cap exposed 3G and 3G-NB usRNAs are targeted primarily for translation. These experiments demonstrate that 5’ cap sequestration, and not DIS exposure or 5’ leader structure itself *per se*, is the major determinant of RNA fate and MDA5-mediated innate immune activation induced by HIV-1 usRNA expression.

We next determined if Rev/CRM1 export was strictly necessary for ^cap^1G usRNA-driven interferon responses, as we and others have previously shown that substitution of Rev/CRM1 RNA export with a retroviral constitutive transport element (CTE) from Mason-Pfizer Monkey Virus (MPMV) fails to induce innate immune responses.(18, 19) To address this, we generated single-round reporter viruses utilizing various RNA export mechanisms with ^cap^1G-only LTRs (**Fig. 6A**). We added either the inactivating M10 mutations in Rev which block Rev/CRM1 export of usRNA (1G-M10) or added M10 in combination with four repeats of CTE (1G-M10/4XCTE) which rescues NXF1-dependent nuclear export of usRNA independent of Rev/CRM1.(29) Upon infection of MDMs, we observed robust infection with all three reporter viruses (**Fig. 6B**). We found that Rev/CRM1 export was strictly required for type I IFN secretion and that 1G-M10/4XCTE infection resulted in only minimal increases in IP-10 secretion relative to 1G-M10 infection (**Fig. 6C/D**). Interestingly, when we probed infected MDM lysates for usRNA abundance via RT-qPCR, we observed low abundance of usRNA in 1G-M10/4XCTE-infected cells, but no defects in p55^Gag^ expression (**Fig. 6E/F**). Altogether, these results support a model where CTE/NXF1-exported ^cap^1G usRNAs are efficiently translated but not sensed by MDA5, similarly to Rev/CRM1-exported ^cap^3G usRNA transcripts (**Fig. 6G**). Collectively, these findings suggest that the immunostimulatory potential of HIV-1 usRNA is modulated by both cap sequestration and nuclear export trafficking itinerary, thus determining its susceptibility to surveillance by MDA5 in the cytoplasm of infected cells.

**Fig. 6:**
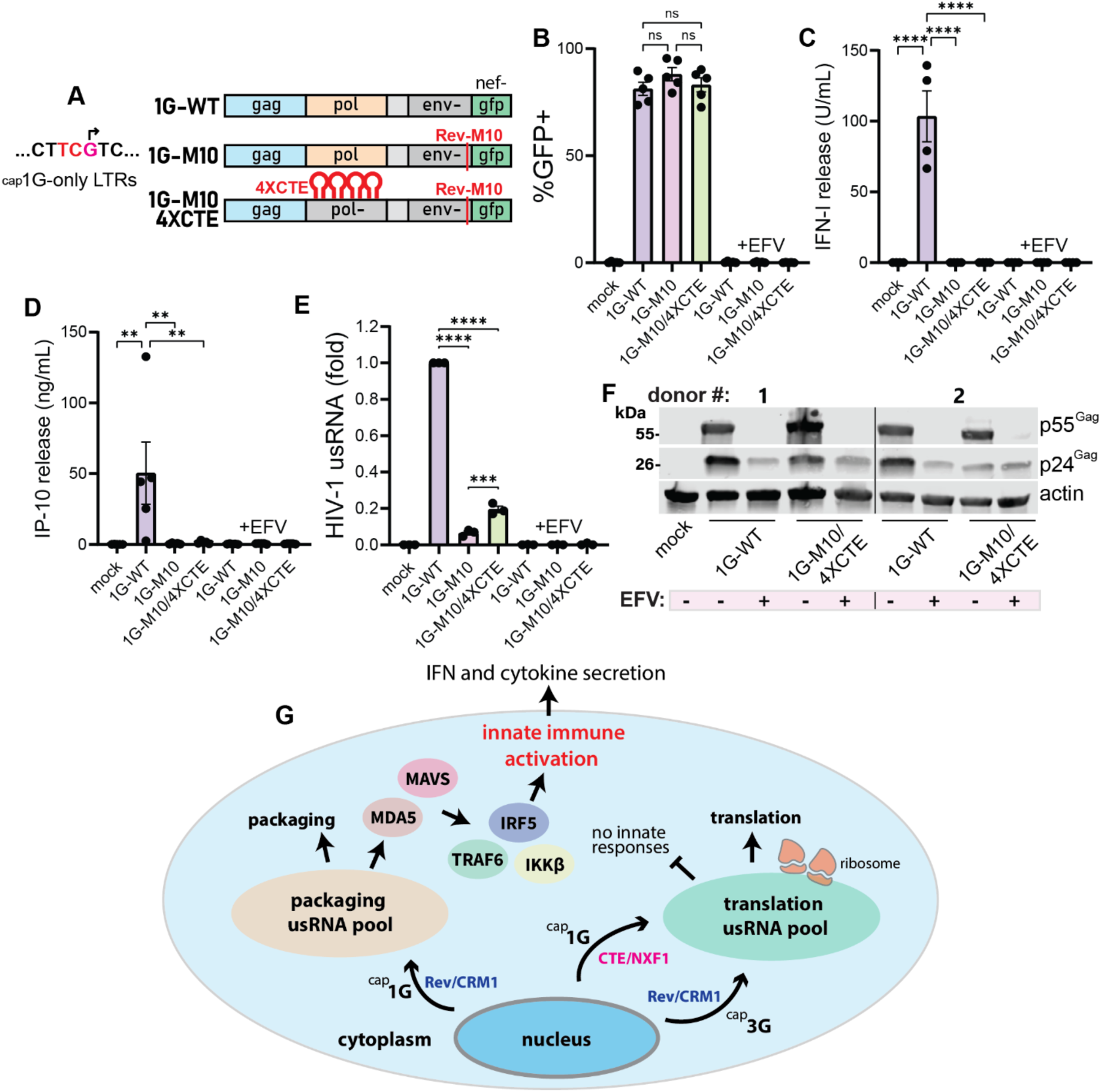
MDA5 sensing of ^cap^1G transcripts requires Rev/CRM1 nuclear export. (**A**) Schematic of HIV-1Δenv/GFP reporter virus genomes cloned into NL4-3 ^cap^1G-only LTR proviral backbone. (**B**-**F**) MDMs were co-infected with VSV-G-pseudotyped reporter viruses and SIV3+ VLPs with or without efavirenz (EFV) treatment. Plots show GFP positivity (**B**), bioactive type I IFN release (**C**), IP-10 release (**D**), relative usRNA abundance (**E**), and Gag expression analyzed by Western blot (**F**) at 3 dpi. (**G**) Model of the findings of this and previous studies, highlighting cytoplasmic pools of HIV-1 usRNA subject to different fates and immunological outcomes. Plots show mean +/- SEM and each data point is an individual primary cell donor. Data was analyzed by one-way ANOVA with Tukey’s post-test, ns: p≥0.05; **: p<0.01; ***: p<0.001; ****: p<0.0001.

## DISCUSSION

The cell-intrinsic innate immune system manages the task of detecting foreign RNAs among a massive background of abundant and diverse host RNA species.(30, 31) In this report, we establish that ^cap^3G unspliced HIV-1 transcripts, which contain an exposed 5’ cap and are efficiently translated utilizing host translational machinery, are remarkably non-immunostimulatory. In contrast, ^cap^1G unspliced transcripts, which contain a sequestered 5’ cap, are recognized by MDA5 to activate innate immune signaling cascades. Since nuclear export and cytoplasmic packaging of ^cap^1G-RNA is an immutable step in the virus life cycle, we hypothesize that TSS heterogeneity to generate ^cap^3G usRNA expression by HIV-1 is a viral innovation that represents viral co-option of immunologically silent mRNA translation pathways to minimize induction of type I IFN responses during viral replication. Importantly, cap sequestration is an essential determinant of efficient viral genome packaging, suggesting that innate immune activation is the putative fitness cost to effectively generate high titers of infectious virus particles.(27) Further, preservation of ^cap^3G unspliced HIV-1 transcript expression in macrophages to facilitate innate immune evasion may at least partially explain the striking conservation of heterogeneous TSS usage among the HIV-1/SIVcpz/SIVgor lineages, as our previous reports have observed only minor replicative defects in ^cap^1G-only virus-infected CD4+ T cells.(22, 32)

We conclude that the innate immune responses observed here are most likely due to usRNA and not singly-spliced transcripts, as splicing of the *gag*/*pol* intron at SD would inhibit the native folded ^cap^1G double three-way junction structure, causing unfolding of the polyA hairpin and making cap sequestration, which triggers detection by MDA5, unlikely. This would agree with a previous report indicating that MDA5 does not engage singly-spliced HIV-1 RNAs.(20) In characterizing viral RNA expression in primary macrophages, we found a greater accumulation of ^cap^1G usRNA levels compared to ^cap^3G in both wild-type and 1G/3G mutant virus infection. In macrophages, because virus particles are frequently retained post-budding in CD169+ virus-containing compartments (VCCs)(33, 34) and particles produced during wild-type or 1G infection are presumably more likely to contain packaged unspliced RNA transcripts than those produced during 3G infection, a higher abundance of ^cap^1G usRNA may not necessarily reflect direct transcriptional differences. However, in line with previous studies, we also found that ^cap^1G RNA expression was biased towards unspliced RNA production.(25) Further, we additionally speculate that cellular turnover of actively translated ^cap^3G transcripts may be faster than ^cap^1G transcripts destined for packaging in macrophages, though the cellular mechanisms postulated to account for differential lifespan and decay of ^cap^3G usRNA remain to be definitively determined. Notably, previous studies have identified translation-dependent destabilization of mRNAs in human cells.(35) While ^cap^3G usRNA abundance was lower than ^cap^1G, we repeatedly did not observe defects in Gag production or particle release in ^cap^3G-only infection, implying that ^cap^3G expression alone is sufficient to drive efficient Gag and Gag-Pol translation while minimizing cytoplasmic usRNA abundance and innate immune activation. Further, we consistently observed a lower ratio of type I IFN production to usRNA expression in 3G infection compared to wild-type or 1G infection, indicating that either the per-transcript immunogenicity of ^cap^3G unspliced transcripts is lower than ^cap^1G, or MDA5 sensing of cytoplasmic HIV-1 usRNA requires a threshold level that ^cap^3G expression does not achieve, or both.

In infections with ^cap^1G-M10/4XCTE virus, where CTE-containing usRNA transcripts access NXF1-mediated rather than CRM1-dependent usRNA nuclear export, we observed an absence of type I IFN production and low usRNA abundance, but robust rescue of *de novo* p55^Gag^ production. We therefore speculate that the relatively small and/or rapidly turned-over pool of cytoplasmic ^cap^1G-M10/4XCTE usRNA transcripts, though etiologically distinct, are functionally similar to ^cap^3G usRNA transcripts and preferentially translated, leading to co-translational decay-dependent usRNA turnover and dampening of innate immune responses. This model would explain previous observations by our group and others that CTE-exported HIV-1 usRNA is not immunostimulatory.(18, 19) In addition, remarkably, this model is consistent with previous reports demonstrating that unspliced murine leukemia virus (MLV) RNA transcripts utilize both CRM1- and NXF1-dependent pathways for nuclear export, and only CRM1-exported MLV-usRNA is effectively packaged into nascent virions, while NXF1-exported MLV-usRNA is instead translated to produce Gag and Gag-Pol polyproteins.(36) Our previous work has also established that CTE-exported HIV-1 usRNA does not effectively compete with Rev/CRM1-exported HIV-1 usRNA for packaging, again implying that NXF1-exported usRNAs have a packaging defect, similar to Rev/CRM1-exported ^cap^3G usRNAs.(37) In summary, the findings in this report and others point to the selection of unspliced retroviral RNA fate and function occurring in the nucleus, which in HIV-1 infection is controlled by transcription start site and determines the immunological outcome of HIV-1 usRNA expression in human macrophages.(36, 38, 39) While HIV-1 exclusively uses Rev/CRM1-mediated nuclear export for unspliced transcripts, ^cap^3G usRNA effectively mimics host mRNA and facilitates robust viral structural/enzymatic polyprotein production while likely minimizing the innate immune cost of expression of ^cap^1G unspliced transcripts required for efficient genome packaging.

Our data additionally point to the sequestration/exposure of the 5’ usRNA cap playing a major role in both the function and immunogenicity of HIV-1 usRNA. A large diversity of positive and negative strand RNA viruses encode mechanisms to both acquire 5’ caps and achieve a host-like methylation pattern on 5’ cap structures on *de novo*-transcribed RNAs, both to achieve efficient translation and to neutralize innate immune responses.(40, 41) Notably, in coronavirus infection, virus-encoded cap 2’-O-methylation activity prevents activation of MDA5 and influences coronavirus pathogenesis in mice.(42, 43) In HIV-1 infection, immunogenicity of cap-sequestered ^cap^1G usRNAs could directly reflect MDA5’s biochemical affinity for uncapped RNAs. Given our previous finding that HIV-1 usRNA-induced innate immune activation requires membrane association,(18) MDA5’s ability to detect cap-sequestered ^cap^1G HIV-1 usRNAs associated with the plasma membrane or coronavirus RNAs lacking cap-methylation associated with double membrane vesicle assembly sites may highlight the proximity of an uncapped or insufficiently-methylated capped RNA to a membrane site as a distinct pathogen-associated molecular pattern (PAMP).(44, 45) Additional variables could theoretically include regulation of MDA5 by the cap-exposed or cap-sequestered usRNA-associated proteome or post-transcriptional RNA modifications.

Alternatively, driven by 5’ cap exposure in ^cap^3G HIV-1 usRNA transcripts, translation itself may repress MDA5 sensing by promoting co-translational decay of usRNAs or obfuscating access to physical RNA-protein interactions with MDA5. Protection of efficiently ribosome-translated RNAs from MDA5 sensing could theoretically decrease the propensity of the innate immune system from spontaneously recognizing host mRNAs that contain significant secondary structure. Further, efficiently-translated Rev/CRM1-exported ^cap^3G usRNAs may enter a pool of cytoplasmic mRNAs not sampled by MDA5, similarly to Rev/CRM1-exported ^cap^3G-NB usRNA transcripts and NXF1-exported transcripts (including multiply-spliced retroviral transcripts and CTE-exported usRNA transcripts), highlighting a probable intracellular localization requirement for MDA5 sensing. In total, our data indicate that HIV-1 usRNA leader structure, function, and intracellular localization are intricately linked to susceptibility to innate immune sensing. Further, these results provide a rationale for HIV-1 dependence on TSS heterogeneity for generating at least two usRNA populations (^cap^1G and ^cap^3G) to dampen MDA5 sensing and overcome restrictions to viral propagation in humans.

## MATERIALS AND METHODS

### Viruses, plasmids, and cloning

The NL4-3, NL4-3 1G, and NL4-3 3G proviral plasmids have been previously described.(22) The NL4-3 3G-NB proviral plasmid has also been previously described.(28) The SIV3+ plasmid used for producing SIV3+ VLPs has been described.(23) The plasmid encoding the G protein of vesicular stomatitis virus (HCMV-G) and the HIV-1 packaging plasmid (psPAX2) used for production of infectious viruses have been described.(18) To generate ^cap^1G-only or ^cap^3G-only single-round GFP-expressing viruses, the proviral genomes of single-round (Δenv, GFP in place of *nef*) Lai-based plasmids (LaiΔenvGFP, LaiΔenvGFP-M10, and LaiΔenvGFP-M10-4XCTE) used in previous studies were excised via *BssHII/XhoI* restriction sites and ligated into the NL4-3 1G or 3G backbones.(18) LaiΔenvGFP and LaiΔenvGFP-M10 themselves were additionally used for smFISH (WT-LTR, WT-LTR/M10). The MDA5-FLAG plasmid used for RNA immunoprecipitation studies was a generous gift from Dr. Kate Fitzgerald (University of Massachusetts Chan Medical School).

All viruses used were pseudotyped with VSV-G envelope protein (regardless of intact or deleted *env*). Viruses were produced via calcium phosphate-mediated transfection of HEK293T cells as described previously.(18) Briefly, HEK293T seeded in 10 cm dishes were transfected with 10-12 μg of plasmids using calcium phosphate/BES-buffered saline. Virus-containing cell supernatants were harvested 48 hours post-transfection, 0.45 μm syringe-filtered and then ultracentrifuged at 100,000 x g in an SW28 rotor for 2 hours at 4°C on a 20% sucrose cushion. The viral pellet was resuspended in DPBS (Gibco, cat. # 14190-144), aliquoted, stored at −80°C, and subsequently titered for infectivity using the TZM-bl assay, as previously described.(18) To produce VSV-G-pseudotyped replication-competent NL4-3-viruses, a plasmid ratio of 10:1 provirus/HCMV-G was utilized. To produce VSV-G-pseudotyped single-round Δenv/GFP viruses, a plasmid ratio of 6:6:1 provirus/psPAX2/HCMV-G was transfected in HEK293T cells. To produce Vpx-containing SIV3+ VLPs, a plasmid ratio of 10:1 SIV3+/HCMV-G was used, and SIV3+ VLPs were titered for p27 content before use with a commercially available p27 ELISA kit (XpressBio, cat. # SK845).

### Cell lines and cell culture

HEK293T cells were acquired from ATCC (cat. # CRL-3216). TZM-bl cells used for titering viruses were acquired from the NIH HIV Reagent Program (cat. # HRP-8129). 293 ISRE-luc cells used for type I IFN bioassay were a generous gift from Drs. Junzhi Wang (National Institute for the Control of Pharmaceutical and Biological Products, China) and Xuguang Li (University of Ottawa). HEK293T, TZM-bl, and 293 ISRE-luc cells were cultured in D10 medium, composed of DMEM (Gibco, cat. # 11965-092) supplemented with 1% Pen-Strep (Gibco, cat. # 15-140-122) and 10% heat-inactivated fetal bovine serum (FBS, Gibco, cat. # A52568-01). MDMs were cultured in R10 medium, made of RPMI1640 medium (Gibco, cat. # 11875-093) supplemented with 1% Pen-Strep and 10% FBS. All cell lines used in these studies were routinely tested for mycoplasma contamination.

### Isolation and differentiation of primary cells

Human peripheral blood mononuclear cells (PBMCs) were isolated from de-identified leukopaks obtained from NY Biologics (Long Island, NY) as described previously.(18) CD14+ cells were isolated from PBMCs using anti-CD14 magnetic beads (Miltenyi Biotech, cat. # 130-050-201) according to the manufacturer’s protocol. To differentiate CD14+ primary monocytes to macrophages, CD14+ cells were plated on low-attachment six well plates (Corning, cat. # 3471) in RPMI media supplemented with 10% heat-inactivated human AB serum (Sigma, cat. # H4522) and 20 ng/mL human M-CSF (Peprotech, cat. # 300-25). After 5-6 days of culture, cells were lifted, counted, and re-plated on normal-attachment plates in R10 medium (or on low-attachment plates if cells were to be analyzed via flow cytometry). CD4+ T cells were isolated from PBMCs using anti-CD4 magnetic beads (Miltenyi Biotech, cat. # 130-045-101) and activated with 2% phytohemagglutinin (PHA, Gibco, cat. # 10576015) and 50 U/mL IL-2 (NIH/NIAID HIV Reference Reagent Program, cat. # 136) for two days before washing in DPBS and resuspension in R10 medium supplemented with 50 U/mL IL-2.

### Infections

MDMs and activated CD4+ T cells were infected with viruses at MOI 1 based on TZM-bl infectious titer. Infections were carried out in the presence of polybrene (Millipore-Sigma, cat. # TR1003G, 10 µg/mL) with or without addition of Vpx-containing SIV3+ VLPs. Cells were spinoculated at 2300 rpm at room temperature for 1 h, incubated at 37°C for 2-3 h, followed by washing with DPBS and culture in fresh R10 medium (with 50 U/mL IL-2 for CD4+ T cells). Cells were harvested at 3 dpi unless otherwise noted. For replication-competent virus infections (MOI 1), 5×10^5^ MDMs were plated in 24-well plates and co-infected with SIV3+ VLPs (5 ng p27^Gag^). For GFP-expressing single-round virus infections, 1×10^6^ MDMs were plated in 12-well plates and co-infected with SIV3+ VLPs (15 ng p27^Gag^).

### Antibodies and drugs

Antibodies used included anti-p24^Gag^ (KC57-RD1, Beckman Coulter cat. # 6604667, mouse monoclonal, used at 1:50 for flow cytometry), anti-p24^Gag^ (NIH/NIAID HIV Reagent Program cat. # HRP-20068, and mouse monoclonal clone # AG3.0, used at 1:500 for co-staining with smFISH). The following antibodies were used for western blot analysis: anti-β-actin (Invitrogen cat. # AM4302, mouse monoclonal, used at 1:5000), anti-β-actin (Sigma-Aldrich cat. # A2066, rabbit polyclonal, used at 1:5000), anti-MDA5 (Proteintech cat. # 21775-1-AP, rabbit polyclonal used at 1:1000), anti-RIG-I (AdipoGen cat. # AG-20B-0009-C100, mouse monoclonal, used at 1:1000), anti-MAVS (Thermo Fisher cat. # PA5-17256, rabbit polyclonal, used at 1:1000), anti-p24^Gag^ (NIH/NIAID HIV Reagent Program, cat. # ARP-6457, mouse monoclonal clone # 24-2, used at 1:1000). For immunoprecipitation, anti-FLAG (Sigma cat. # F3165), and mouse IgG (Novus cat. # NBP1-97019) were used. Secondary antibodies used were goat anti-mouse DyLight 680 (for western blot analysis, Invitrogen cat. # 35518, used at 1:10,000) goat anti-rabbit DyLight 800 (for western blot analysis, Invitrogen cat. # SA5-35571, used at 1:10,000), goat anti-mouse AlexaFluor647 (for anti-p24^Gag^ co-staining with smFISH, Thermo Fisher, cat. # A-21237, used at 1:200).

Antiretrovirals and drugs used included lenacapavir (1 µM, NIH HIV Reagent Program, cat. # HRP-20266), efavirenz (1 µM, NIH HIV Reagent Program, cat. # HRP-4624), raltegravir (30 µM, Selleckchem, cat. # S2005), KPT-330 (1 µM, Selleckchem, cat. # S7252), lopinavir (1 µM, NIH HIV Reagent Program, cat. # HRP-9481).

### siRNA transfection

Pooled siRNA stocks were obtained from Horizon Discovery (cat. # MDA5: L-013041-00-0005, RIG-I: L-012511-00-0005, MAVS: L-024237-00-0005, control: D-001810-10-05). siRNA was transfected into MDMs two days prior to infection using TransIT-X2 (Mirus, cat. # 6000) at 50 nM final siRNA concentration. To assay for knockdown efficiency, transfected MDMs were washed and either lysed for Western blot analysis on the day of infection in RIPA buffer or lysed for RT-qPCR in Qiagen RLT buffer on the day of infection or at 3 dpi.

### CaDAL assay

5’ ends of unspliced HIV-1 RNAs were analyzed by cap-dependent ligation/PCR assay as previously described.(22) Total RNA from infected MDMs was analyzed via cDNA synthesis from an RT primer downstream of the major splice donor site to restrict analysis to solely unspliced RNAs. ^cap^1G/^cap^3G ratio within a given sample is quantitative, while comparisons between samples or cell types are not quantitative.

### RT-qPCR

RT-qPCR was performed as previously described.(18) Briefly, total RNA was isolated using a Qiagen RNEasy kit (cat. # 74106). cDNA was generated from equivalent amounts of total RNA using oligo-dT primers (Invitrogen, cat. # 18418020) and SuperScript III RT (Invitrogen, cat. # 18080051). qPCR was performed with 5-20 ng cDNA using 2X Maxima SYBR Green master mix (Thermo Scientific, cat. # FERK0242) and primer sets outlined in **Table S1**. Quantification was performed using the ΔΔCt method with GAPDH used as an internal control.

### Digital droplet PCR

To analyze splicing patterns between NL4-3/1G/3G viruses, RT-ddPCR was performed similarly to previous descriptions with modifications.(24) cDNA was generated from 50 ng total RNA from infected MDMs as described above, using random hexamers as primers. cDNA was diluted 1:1500 (or no-RT control samples were diluted 1:150) and mixed with Supermix for Probes (no dUTP) (Bio-Rad, cat. # 186-3023) and primer/probe sets targeting LTR, *nef*, *env*, and *gag* outlined in **Table S1**. Droplets were generated using a Bio-Rad QX200 ddPCR droplet generator. LTR and *nef* probes were read in the FAM channel (with different probe concentrations to distinguish the signals), while *env* and *gag* were read in the HEX channel (with different probe concentrations to distinguish the signals). The final concentration of primers/probes in droplets was: 900 nM (all primers), 250 nM (LTR-FAM and *gag*-HEX probes), and 150 nM (*nef*-FAM and *env*-HEX probes). PCR was performed and droplets were read using a Bio-Rad QX200 droplet reader. Data was analyzed in QuantaSoft software, and gates were drawn to distinguish the LTR, *nef*, *env*, and *gag* signals. To analyze splicing patterns, only LTR/nef, LTR/env/nef, and LTR/gag/env/nef droplets were compared (all other droplet populations were discarded) and relative transcript abundance was calculated for each primary cell donor.

### smRNA-FISH

MDMs were cultured on gelatin-coated glass coverslips at 3×10^5^ cells/well in 24-well plates before infection with LaiΔenvGFP (WT-LTR) or LaiΔenvGFP with NL4-3 LTR/backbone sequences (1G-LTR or 3G-LTR) at MOI 0.5 with SIV3+ VLP co-infection (1 ng p27 per well). Infected MDMs were cultured for 3 days before fixation with formaldehyde and permeabilization with DPBS/Tween-20 (0.1%, nuclease-free, Promega, cat. # PRH5152). Coverslips were then blocked with 1% bovine serum albumin (nuclease-free, Sigma-Aldrich, cat. # 126615-25ML) and stained for p24^Gag^. Alternatively, coverslips were stained with wheat germ agglutinin conjugated to AlexaFluorPlus568 (WGA, 1 µg/mL for 10 min on ice, Invitrogen, cat. # W56133) to label plasma membrane before fixation and blocking. After anti-p24^Gag^ immunostaining, coverslips were fixed again with formaldehyde before smFISH.

To perform smFISH, stained coverslips were equilibrated with wash buffer A (Stellaris, cat. # SMF-WA1-60) before hybridization in hybridization buffer (Stellaris, cat. # SMF-HB1-10) containing 125 nM probes. De-ionized formamide (Millipore Sigma, cat. # 4610-100ML) and nuclease-free water were added to wash and hybridization buffers before use per the manufacturer’s instructions. For p24^Gag^ staining, probes consisted of 48 CALFluor590-labeled oligos designed against HIV-1 Lai *gag/pol* as outlined in **Table S2** (Stellaris). For WGA-568 staining, probes consisted of 48 Quasar670-labeled oligos also outlined in **Table S2**. Coverslips were hybridized overnight at 37°C in a humidified chamber. Coverslips were then washed with wash buffer A at 37°C for 30 min, followed by DAPI staining at 37°C or 30 min (5 ng/mL in wash buffer A), and then equilibrated with wash buffer B (Stellaris, cat. # SMF-WB1-20) for 5 mins at room temperature before mounting onto a slide using VECTASHIELD mounting medium (Vector Laboratories, cat. # H-1900-10).

Images were taken using a Leica SP5 confocal microscope. For colocalization analysis, regions of interest were drawn manually around cells and fluorescent co-localization was quantified using the JACoP plugin for ImageJ/Fiji.

### RNA immunoprecipitation

RNA immunoprecipitation in infected HEK293T cells transfected with MDA5-FLAG was performed as described previously.(21) After UV-crosslinking and cytosolic fractionation followed by immunoprecipitation, RNA was extracted from antibody-coated beads using TRIzol/chloroform extraction before cDNA synthesis, DNase treatment, and qPCR. Primers used for qPCR are listed in **Table S1**. Beads were washed in RIPA low-salt (150 mM NaCl) followed by RIPA high-salt buffer (1M NaCl) before RNA extraction or extraction for Western blot analysis. For Western extraction, beads were boiled with neat 6X loading buffer (containing 4% SDS and 2% β-mercaptoethanol) before bead removal and gel loading. Data is reported as fold enrichment of RT-qPCR signal between anti-FLAG and IgG control immunoprecipitation conditions.

### Type I interferon bioassay

Type I interferon bioactivity was measured using 293 ISRE-luc cells similar to previous descriptions.(18) 4×10^4^ cells were plated with 100 μL of culture supernatant or IFNα2 standard diluted in R10 medium (1.6-200 U/mL, PBL Assay Science cat. # 11100-1) in 96-well plates and incubated at 37°C for 24h. Cells were washed in DPBS, lysed in Glo Lysis buffer (Promega, cat. # E266A), and luciferase activity in cell lysates was determined using Bright-Glo reagent (Promega, cat. # E2610) using a LUMIstar Omega reader (BMG LabTech). For quantification, the lowest standard (1.6 U/mL) was considered the limit of detection and samples with lower luminescence than the limit of detection were reported as 0 U/mL.

### ELISA

IP-10 secreted into culture supernatants was quantified with ELISA (BD, cat. # 550926) after inactivation in DPBS/10% NCS/0.5%TX-100. p24^Gag^ in culture supernatants was detected as previously described using an in-house ELISA.(18)

### Western blot analysis

To perform Western blot analysis, 30 μg of protein (quantified with Bradford assay) was diluted to equivalent volume in DPBS/loading buffer (containing SDS and β-mercaptoethanol), boiled, and loaded into Mini-PROTEAN TGX Precast Protein Gels (BioRad, 4-20% gradient, cat. # 4561094) along with a protein ladder (PageRuler Prestained NIR Protein Ladder, Thermo Scientific, cat. # 26635). Blots were blocked with LI-COR Intercept blocking buffer (LI-COR, cat. # NC1660556) before staining and imaging on a LI-COR Odyssey CLx imager.

### Flow cytometry

To assess infection efficiency with replication-competent viruses, MDMs were seeded on low-attachment plates and lifted using Cellstripper (Corning, cat. # 25-056-CI). Cells were moved to a U- or V-bottom plate, fixed in 4% paraformaldehyde for 15-30 min at 4°C, permeabilized with Perm/Wash buffer (BD, cat. # 554723), and stained with anti-p24^Gag^ antibody for 30 min at room temperature. Flow cytometry was performed immediately using a Cytek Aurora spectral analyzer. Data was analyzed and spectral plots were generated in FlowJo 10.8.1.

### Statistics

Statistics were performed in GraphPad Prism 10.2.3. Two-sample comparisons were analyzed by t-test. For multi-sample comparisons, one-way or two-way ANOVA were performed with Dunnett’s or Tukey’s post-test. For MDMs, each data point represents an individual primary cell donor, while for cell lines, each data point represents an independent experiment. Plots represent the mean +/- SEM of the data, ns: p≥0.05; *: p<0.05; **: p<0.01; ***: p<0.001; ****: p<0.0001.

## Data Availability

The authors declare that the data that support the findings of this study are available from the corresponding author upon reasonable request.

## Acknowledgements

The authors thank the BU-CAMED Flow Cytometry Core Facility for technical assistance and the NIH/NIAID HIV Reagent Program and associated investigators for reagents used in this study. The authors thank NIH for funding used to perform this study. This work was supported by grants R01DA059952 (S.G. and A.H.), R01AI187175 (S.G. and A.H.), R01DA055488 (S.G. and A.H.), R01DA051889 (S.G.), and 5P30AI042853 (S.G., A.H., and H.A.).

## Author contributions

I.H., S.K., A.H., H.A., A.T., and S.G. conceived of and designed the experiments. I.H. and S.G. wrote the manuscript. I.H., H.A., A.H., S.K., A.T. and S.G. edited the manuscript. I.H., S.K., S.R., S.J., J.H., and X.H. performed the experiments.

---

**Fig. S1:**
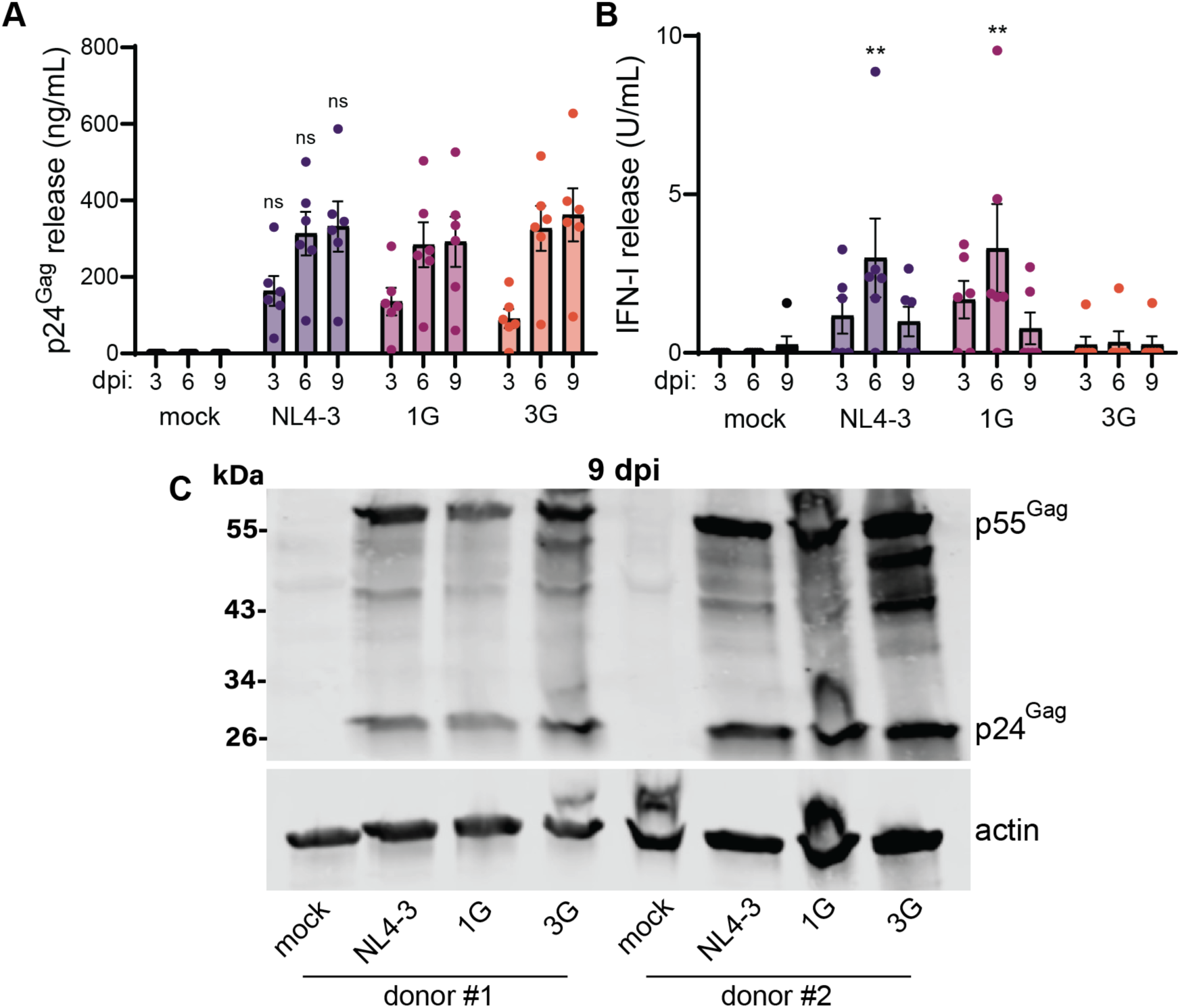
^cap^1G unspliced HIV-1 RNA expression stimulates type I interferon release in macrophages. MDMs were infected with HIV-1 alone (no SIV3+ VLP addition) and culture supernatants were analyzed at 3, 6, or 9 days post-infection. Plots show p24^Gag^ release (**A**) and IFN-I release (**B**) at the indicated timepoint. (**C**) Western blot analyzing Gag expression, harvested at 9 dpi from cells used in (**A**/**B**). Data points are individual primary cell donors, plots show mean +/- SEM. Data was analyzed by two-way ANOVA with Tukey’s post-test (**A**) or Dunnett’s post-test with statistics referring to comparison with mock infection (**B**). (**A**) All virus infection comparisons nonsignificant, comparing the same timepoint across virus infections. (**B**) **: p<0.01 comparing NL4-3 and 1G infection to mock infection at 6 dpi, all other comparisons nonsignificant (p≥0.05). (**C**) Western blot was performed with two individual primary cell donors.

**Fig. S2:**
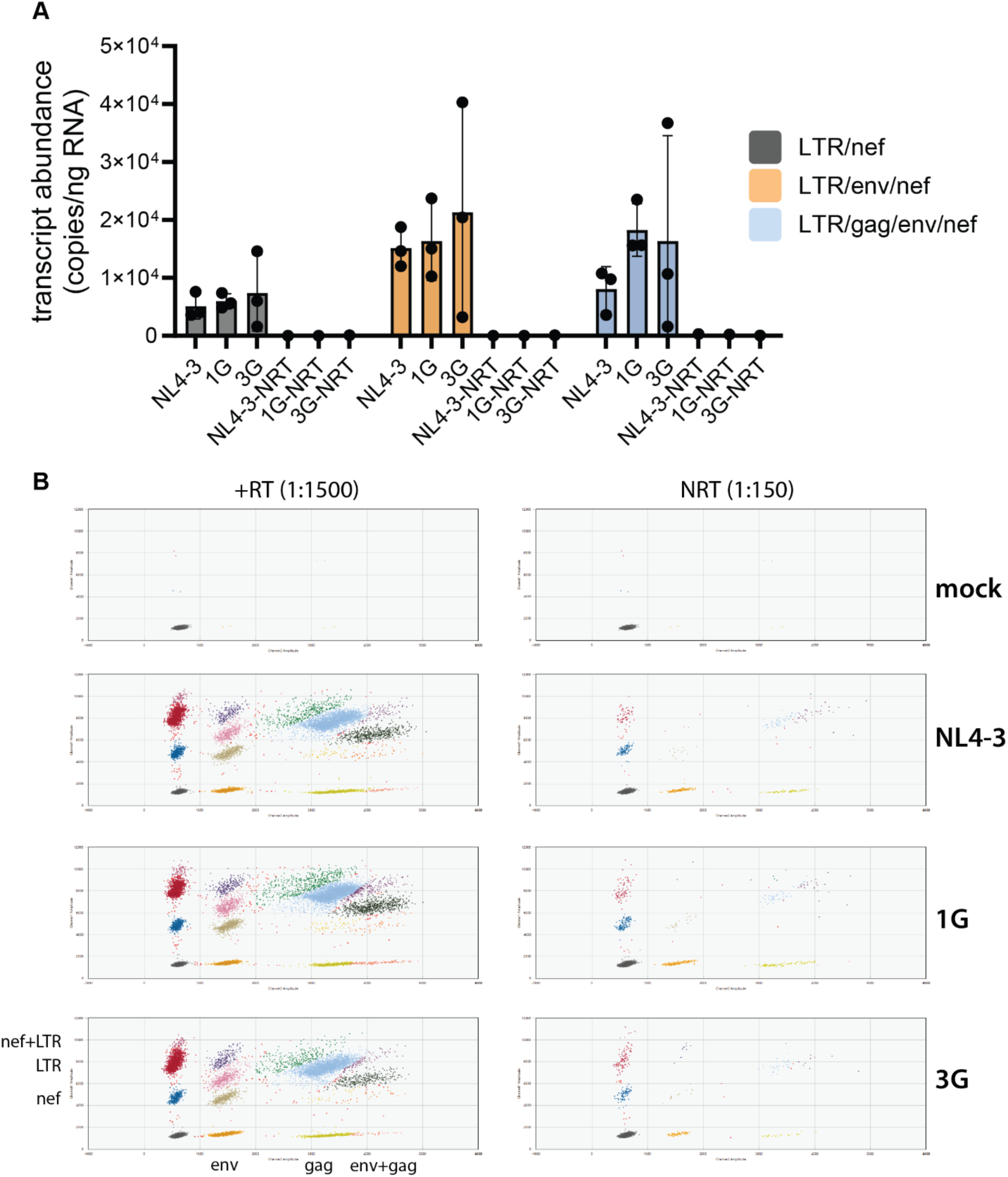
ddPCR for viral transcriptomic analysis in primary macrophages. Additional data from experiment described in Fig. 4C. (**A**) Raw transcript abundance values obtained from ddPCR, NRT: no-RT control. (**B**) Sample gating strategy to distinguish viral RNA populations, for +RT samples diluted 1:1500 or NRT controls diluted 1:150. (**A**) Plot is shown as mean +/- SEM, each data point is an individual primary cell donor. (**B**) Data is from one primary cell donor.

**Fig. S3:**
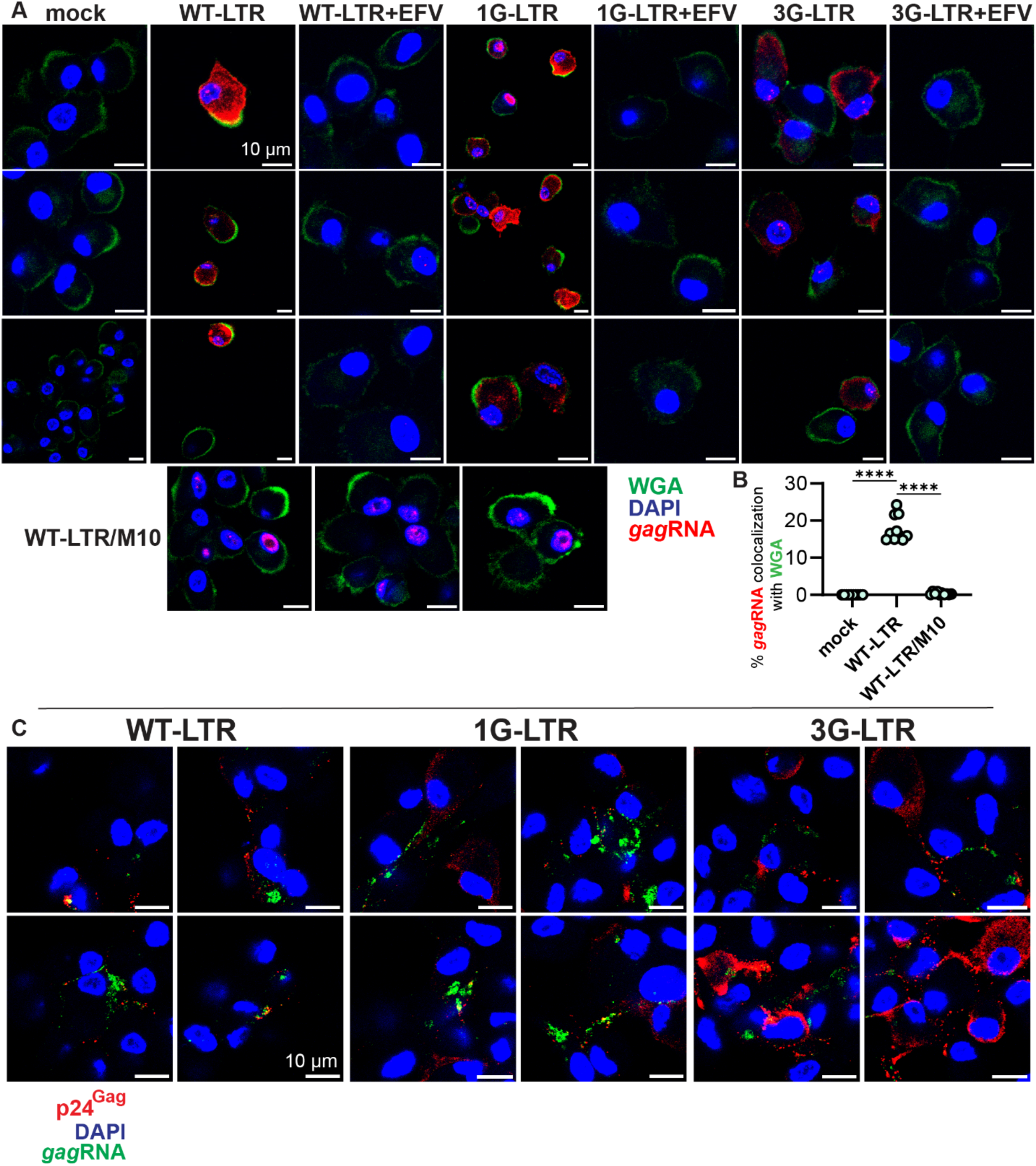
smFISH of ^cap^1G/3G usRNA expression in primary macrophages. Additional data from experiment described in Fig. 4D-**G**. Additional magnified images and colocalization quantification showing smRNA-FISH analysis merged with WGA (**A**, wheat germ agglutinin, plasma membrane stain) or anti-p24^Gag^ co-stain (**C**). In (**B**), % of *gag* RNA signal colocalizing with WGA is shown. EFV: reverse transcriptase inhibitor efavirenz treatment. WT-LTR/M10: wild-type ^cap^1G/3G-expressing virus with inactivating M10 mutation in Rev. Means +/- SEM of individual fields of view from one representative experiment are shown, with data analyzed by one-way ANOVA with Tukey’s post-test, ****: p<0.0001.

**Fig. S4:**
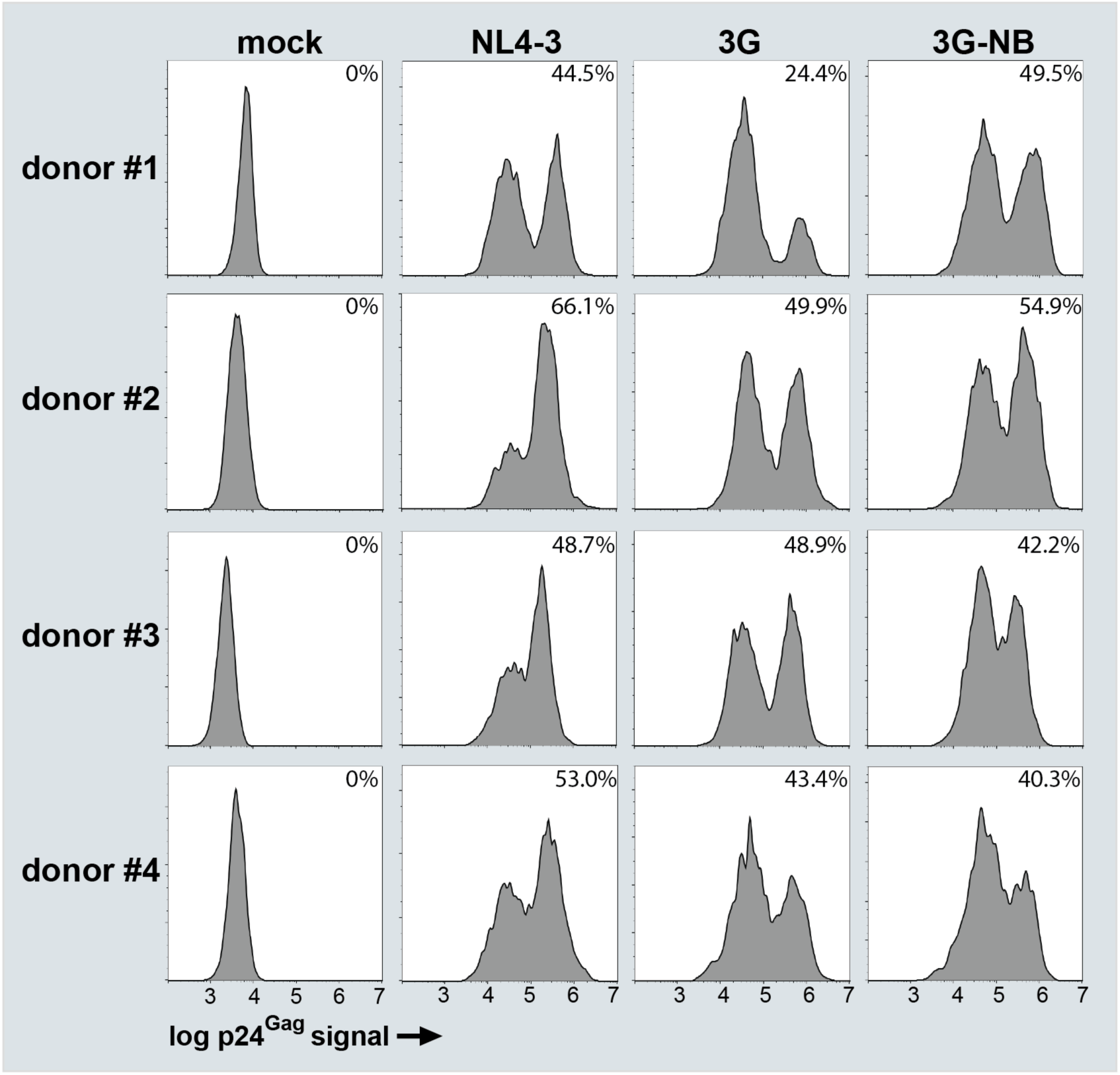
p24^Gag^ production efficiency in 3G and 3G-NB infection. Additional data from experiment described in Fig. 5B. (**A**) Plots show log p24^Gag^ flow cytometry signal as a histogram with positive population quantification for four primary cell donors.

**Table S1:**
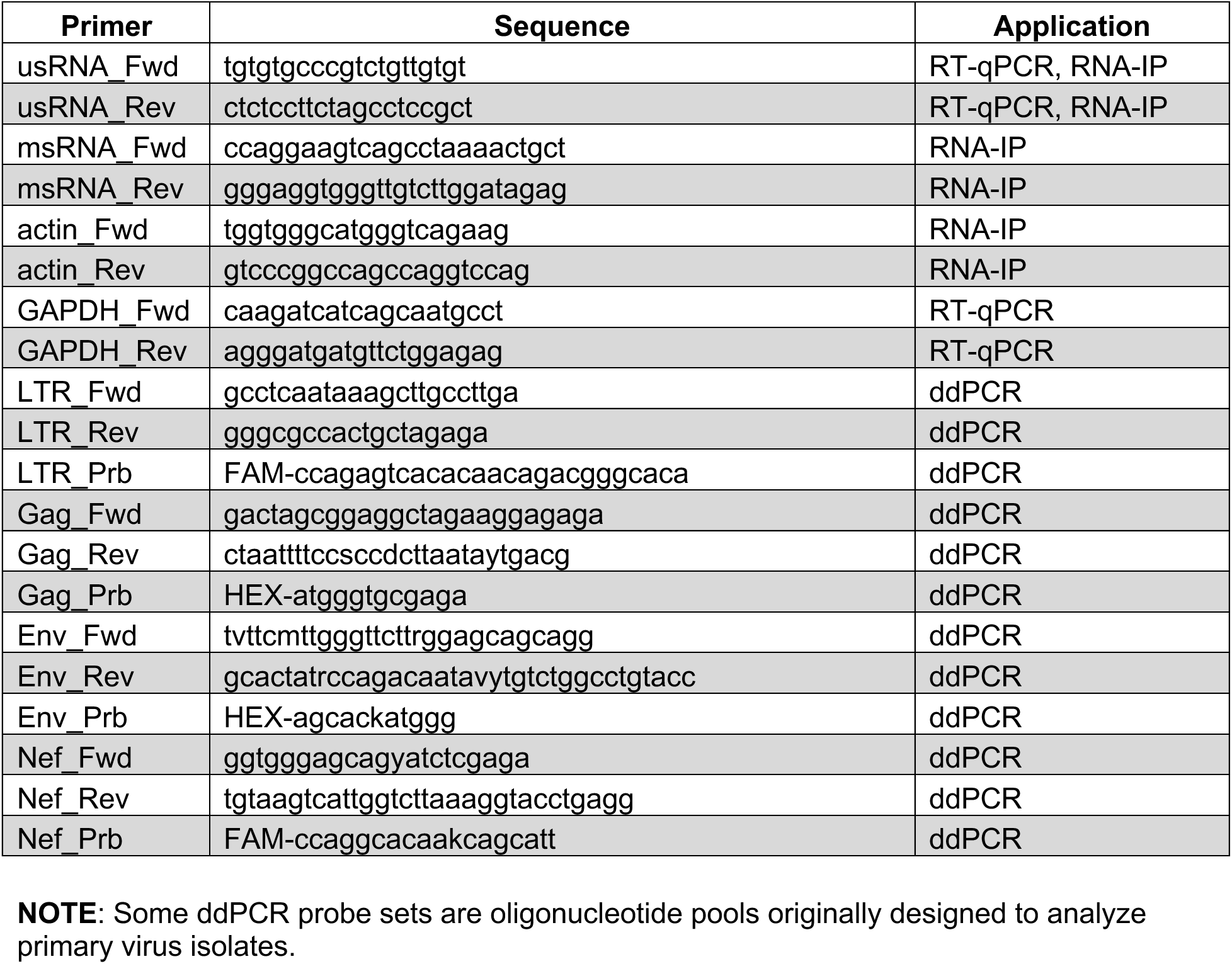
Primers used for PCR-based assays.

**Table S2:**
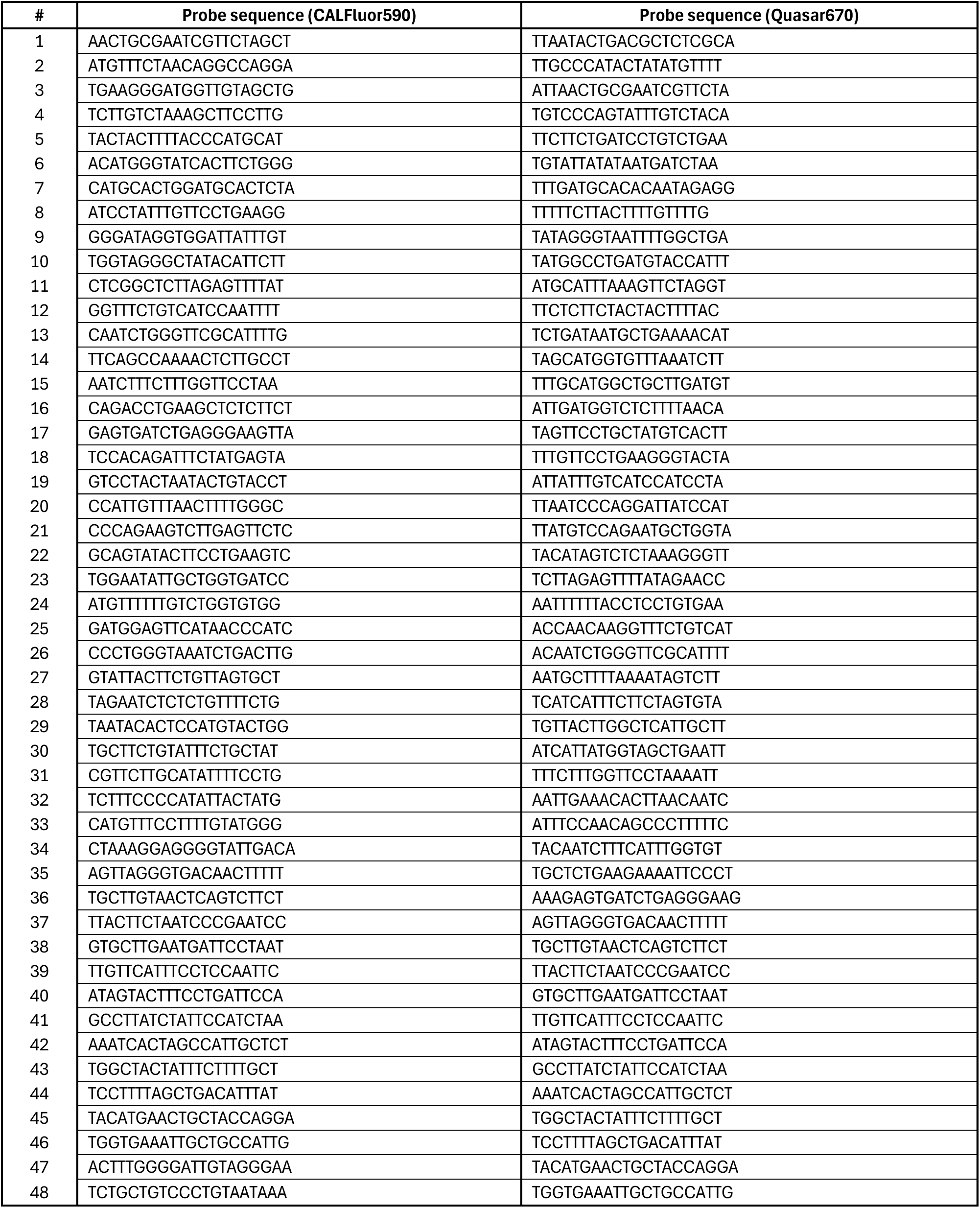
Fluorescent smRNA-FISH probe sequences.

